# Association mapping reveals the role of mutation-selection balance in the maintenance of genomic variation for gene expression

**DOI:** 10.1101/015743

**Authors:** Emily B. Josephs, Young Wha Lee, John R. Stinchcombe, Stephen I. Wright

## Abstract

The evolutionary forces that maintain genetic variation for quantitative traits within populations remain poorly understood. One hypothesis suggests that variation is maintained by a balance between new mutations and their removal by selection and drift. Theory predicts that this mutation-selection balance will result in an excess of low-frequency variants and a negative correlation between minor allele frequency and selection coefficients. Here, we test these predictions using the genetic loci associated with total expression variation (eQTLs) and allele-specific expression variation (aseQTLs) mapped within a single population of the plant *Capsella grandiflora.* In addition to finding eQTLs and aseQTLs for a large fraction of genes, we show that alleles at these loci are rarer than expected and exhibit a negative correlation between phenotypic effect size and frequency. Overall, our results show that the distribution of frequencies and effect sizes of the loci responsible for local expression variation within a single, outcrossing population are consistent with mutation-selection balance.

**Significance:** Biologists have long sought to explain why we see genetic variation for traits in populations despite the expectation that selection will remove most variation. We address this question by using gene expression as a model trait and identifying the genetic loci that affect gene expression in a single, large population of the plant *Capsella grandiflora.* Alleles at loci that affect expression were rarer than expected by chance and there was a negative correlation between phenotypic effect size and frequency of these alleles. These observations are consistent with mutation-selection balance, the hypothesis that variation is maintained by a balance between new mutations and their removal by selection and drift.

## Introduction

Genetic variation for quantitative traits persists within populations despite the expectation that prevalent stabilizing selection will reduce genetic variance. One hypothesis suggests that variation is maintained by a balance between new mutations and their removal by selection and drift, resulting in an excess of low-frequency variants and a negative correlation between minor allele frequency and selection coefficients (1). While studies of allele frequency spectra show that purifying selection is often prevalent in genomic sequence (2–4), little is known about how the genetic variants under selection relate to phenotype, and ultimately, how phenotypic variation is maintained within populations. Association mapping can identify specific loci influencing phenotype providing candidates for further analysis of selection (5). In particular, mapping the local regulatory variants that affect gene expression can identify a large number of genetic loci that affect phenotype. Additionally, mapping the genetic basis of gene expression will answer questions about the basic biology of gene regulation, for example, by testing predictions that conserved non-coding sequences (‘CNSs’) are constrained because they have regulatory function (6).

Early eQTL studies mapped expression divergence between two lines, finding that many genes have local expression QTL (7, 8). These studies have provided insight into selection on eQTLs; for example, a correlation between recombination rate and eQTL density implies that background selection is a dominant force acting on expression variation in *Caenorhabditis elegans* (9) and a skew towards rare allele frequencies in promoters of genes with eQTLs suggests that purifying selection may act on expression variation (10). However, eQTL studies of population-level genetic variation have thus far been limited to a few study systems (11–15) and only one study, in humans, has identified a negative correlation between phenotypic effect size and frequency (14). In addition, human eQTL studies have shown that loci expected to be involved in selective sweeps are more likely to be eQTLs than other loci (16), allele frequencies of eQTLs that increase expression of a potentially deleterious coding SNP are under stronger purifying selection than those that do not (17), and eQTL allele frequencies within populations are linked to local adaptation (18, 19). To date, eQTL studies in plants have used genetic crosses (20–22) or species-wide samples (23–25), making it difficult to distinguish evolutionary forces acting within and between populations. In sum, we currently lack comprehensive tests of selection on within-population eQTLs in any system, especially in plants.

Here, we map local regulatory loci affecting expression in 99 members of a single large population of *Capsella grandiflora* (Brassicaceae), an obligate outcrosser. As might be expected from its large N_e_ and relative lack of population structure, purifying and positive selection are strong in *C. grandiflora*(3, 26), making it an ideal system for investigating the maintenance of genetic variation in the face of selection

## Results and Discussion

We sequenced 22,895,738,517 100bp paired-end reads of DNA from 188 individuals, with a median of 119,321,591 reads per individual. Of these reads, a median of 93% mapped per individual (range: 51%-93%, the two individuals with <80% were not sampled for RNAseq). We called 9,526,786 SNPs with a mean depth of 45 reads per individual. Linkage disequilibrium between SNPs decays rapidly: mean R^2^ between SNPs less than 10bp apart is 0.25, and this decays to 0.12 within 100bp (Fig. S1). An analysis of population structure (27) found that the maximum likelihood number of populations was K=1, suggesting no widespread structure. We measured genome-wide gene expression in 99 of these individuals using RNAseq from young leaf tissue, generating 4,988,540,400 100bp paired-end RNAseq reads with a median of 49,549,336 reads per individual (range: 42,627,096–106,283,910). Of these, a median of 94% (range: 89–95%) mapped to genes (Table S1).

We mapped eQTLs by performing Mann-Whitney U tests comparing expression between individuals homozygous for the most common allele at a given SNP and those heterozygous at that SNP, for all SNPs within 5kb of the transcription start and end sites (Fig. 1). We omitted rare homozygotes from the analysis because most local regulation acts additively in *cis* (12) and low sample sizes for rare variants reduce power. Out of 5,507,316 SNPs tested against the expression of 18,692 genes, 39,628 SNPS are significantly associated with expression of 6,624 nearby genes (FDR = 0.1, p < 8.2 × 10^−4^, Fig. S2 A). These SNPs often clustered locally (Fig. S3 A,B), as would be expected if non-causal SNPs are in linkage disequilibrium with causal SNPs. Patterns of functional enrichment in human eQTLS suggest that SNPs most strongly associated with expression are more likely causal than those showing weaker associations(13), so to prevent variation in linkage disequilibrium from affecting subsequent analyses while increasing the likelihood of retaining causal SNPs, we chose the most significantly associated SNP for each gene for further analysis (N = 6,624). While there are likely multiple causal eQTLs for many genes, choosing one significant SNP per gene allows us to generate a large independent sample of eQTLs for further analysis.

**Figure 1:**
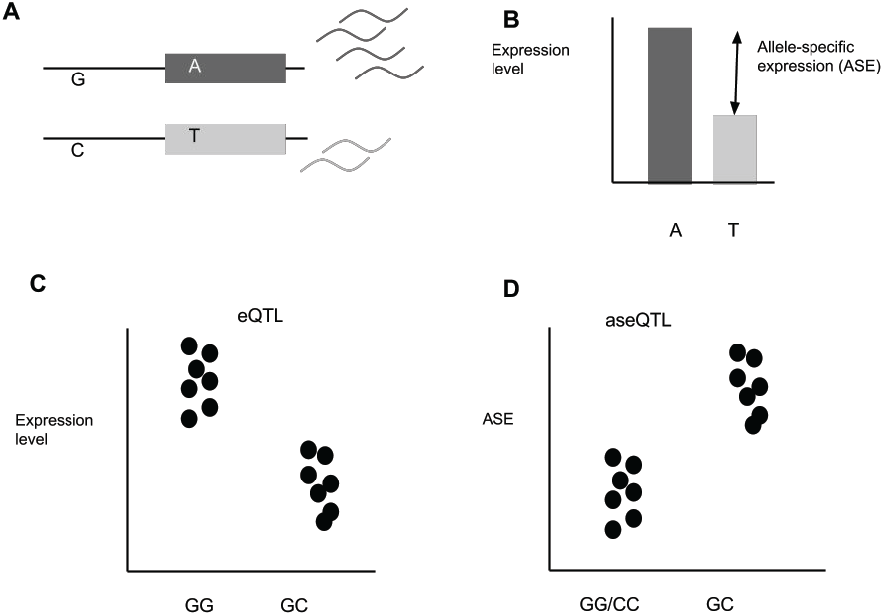
Detecting eQTLs and aseQTLs. (a) A gene model for an individual that is heterozygous at a regulatory locus (G/T) and at an informative coding site (A/T). The G allele increases expression relative to the C allele, (b) causing increased allelic expression of the reads carrying the A allele at the informative heterozygous site. We refer to this difference in allelic expression as “ASE”. (c) eQTLs are detected when there is a significant difference in total gene expression between individuals (represented by black circles) that are homozygous for the common allele of a SNP and individuals that are heterozygous at that SNP. (d) aseQTLs are detected when there is a significant difference in ASE between individuals that are heterozygous at a SNP and homozygous for either allele at that SNP.

If eQTLs act in *cis,* heterozygous eQTLs will cause allele-specific expression (ASE), providing an additional signature of regulatory variation. We measured ASE within individuals by calculating the mean expression difference between alleles, standardized for sequencing depth. We then mapped QTLs for ASE (‘aseQTLs’) by performing Mann-Whitney U tests comparing ASE in individuals that were homozygous at a local SNP and those that were heterozygous at that SNP (Fig. 1). We excluded coding SNPs from this analysis because their genotype might confound ASE measurement. Out of 3,966,423 SNPS tested, 26,957 SNPs were significantly associated with ASE of 5,882 nearby genes (FDR = 0.1, p < 5.4 × 10^−4^, Fig S1 B). Our analysis did not require a directional effect of SNP genotype on ASE, but 22,436 (83%) of the noncoding SNPs associated with ASE have higher ASE in heterozygotes, as would be expected if these SNPs control expression in *cis.* We selected the most strongly associated noncoding SNP per gene for further analysis and we also required that ASE had to be higher in heterozygotes at that SNP than homozygotes, leaving 4,580 aseQTLs (Fig S2 B).

SNPs located near the transcription start site (TSS) and in 5’ UTRs were more likely to be eQTLs and aseQTLs than SNPs further away from the gene (Fig 2A), consistent with data from humans and *Drosophila* (11, 12, 14). In addition, CNSs near the TSS were enriched for eQTLs and aseQTLs relative to non-conserved sites (Fig. 2A), suggesting that genetic variation within CNSs represents a major source of standing variation in gene expression, although bootstrapped confidence limits for these overlap slightly in aseQTLs. In contrast, CNSs in 5’UTRs were not enriched for eQTLs or aseQTLs, consistent with observations that selection strength is relatively similar in conserved and non-conserved sites in these regions(3). However, the detection of a large number of eQTLs outside of conserved regions suggests that regulatory element turnover is common in Brassicaceae (Table S2). There were 2,236 genes that had both eQTLs and aseQTLs, significantly more than expected by chance (X^2^ = 471, p < 2.2×10^−16^). Of these 2,236 genes, 411 had the same SNP most significantly associated both with expression and ASE.

**Figure 2:**
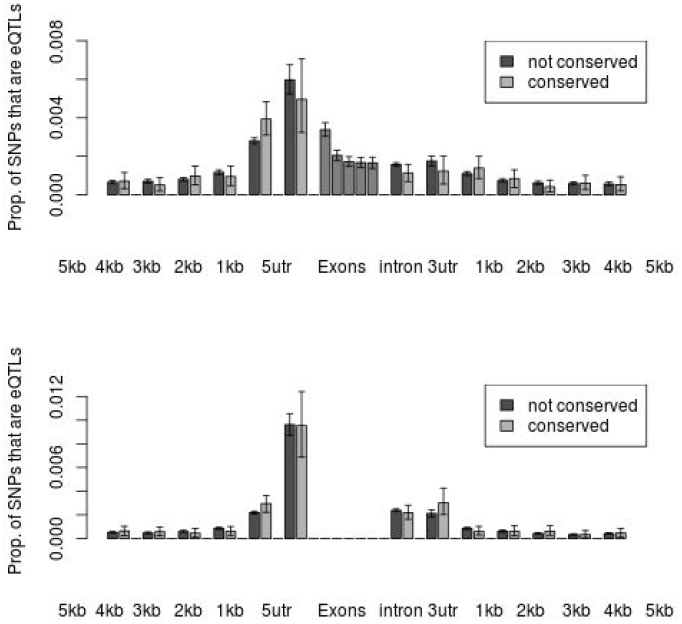
eQTL and aseQTL enrichments by site type. The proportion of SNPs tested in each category that were found to be eQTLs is plotted on the y axis for (a) eQTLs and (b) aseQTLs. The exonic classes were determined by splitting the coding sequence of each gene into 5 equally sized pieces. Note that there were no exonic SNPs included in the aseQTL analysis. Error bars show the 95% confidence intervals from bootstrapping.

Next, we tested eQTLs and aseQTLs for signatures of selection. Purifying selection will reduce the frequency of causal alleles at QTLs, but allele frequency also controls sample size in association studies, affecting QTL detection. Rare alleles have an increased likelihood of false negatives, because of lower power, and false positives, since expression is not normally distributed and an outlier in a small sample is more likely to lead to a positive association than an outlier in a large sample. The increased likelihood of false positives in rare alleles makes evolutionary inferences especially challenging because it mimics the signal of purifying selection.

To generate an appropriate null distribution for QTL allele frequency, we permuted assignments between expression level and genotype for every gene 1000 times and ran eQTL analyses using permuted data. On average, 3,258 SNPs were associated with total expression in our permutations, consistent with an FDR of 0.1, since 39,628 SNPs were associated with the observed data. However, observed eQTLs from un-permuted data were significantly rarer than those found in permuted data (mean N=2,047), consistent with the action of purifying selection (Fig. 3A). This observation is conservative, because we have not accounted for reduced power to detect associations on rare alleles. We also investigated permuted aseQTLs, and found on average 3,194 SNPs associated with ASE in each permutation, which is slightly more than expected given our FDR of 10% (26,597 SNPs were associated with ASE in un-permuted data). As with eQTLs, aseQTLs were significantly rarer than those found in permuted data (Fig. 3B). These results hold when we designate a random significantly-associated SNP per gene as the eQTL or aseQTL (Fig S4 A,B). In addition, eQTLs and aseQTLs are significantly rarer than permuted eQTLs and aseQTLs when only SNPs 1–5 kb upstream or downstream of genes are considered (Figure S5, A,B), and when sites are separated into high and low recombination sets or by substitution type (Figure S5 C,D). Thus, the frequency distribution of both eQTLs and aseQTLs is consistent with the predominance of mutation-selection balance.

**Figure 3:**
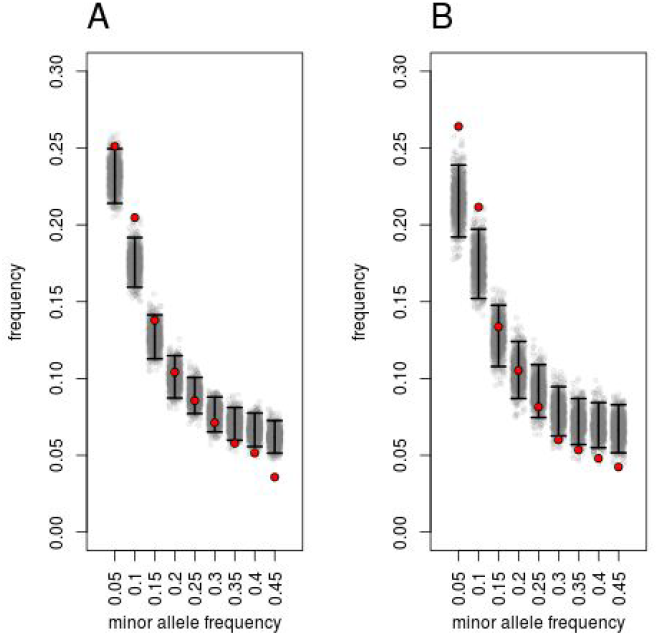
The site frequency spectra of eQTLs and aseQTLs. Minor allele frequencies of (a) eQTLs and (b) aseQTLs for observed data (red circles) and permuted data (gray circles, black lines are 95% confidence intervals).

**Figure 4:**
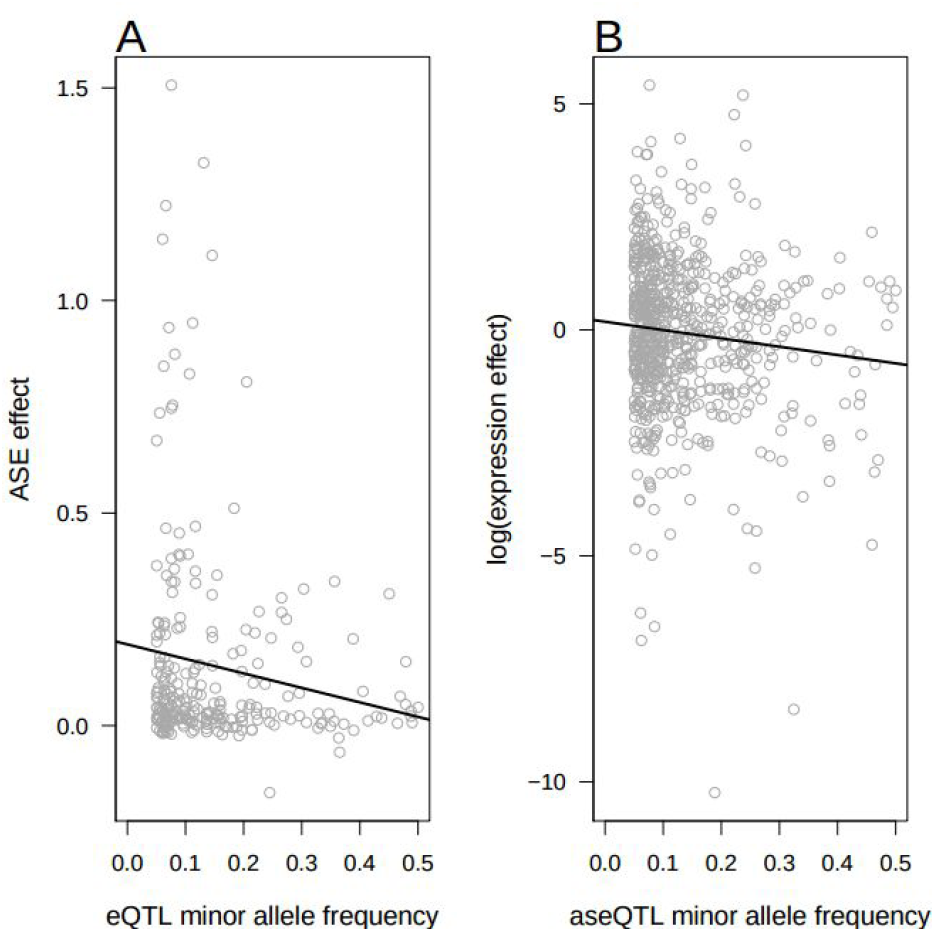
The relationship between minor allele frequency and effect size. (a) eQTL minor allele frequency is plotted against the effect of that SNP on ASE, calculated as the mean difference in ASE between individuals heterozygous at the eQTL and individuals homozygous at the eQTL. Negative values occur when the the homozygote for the eQTL has greater ASE than the heterozygote. The trend line is calculated by linear regression (b) aseQTL minor allele frequency plotted against the effect of the aseQTL on total gene expression, calculated by taking the log of the absolute value of the mean difference in expression between individuals heterozygous at the aseQTL and individuals homozygous for the common allele at the aseQTL. The trend line was calculated by regression between minor allele frequency and the log of the expression effect.

We incorporated effect sizes to test for an additional signature of selection. Theory predicts that mutation-selection balance will maintain mutations at frequencies inversely proportional to the strength of selection acting against them (1), suggesting that QTLs under purifying selection should show a negative correlation between minor allele frequency and phenotypic effect size, assuming that phenotypic effect size correlates with the strength of selection. However, this correlation is also expected if QTLs evolve neutrally for two reasons. First, we have low power to detect rare small-effect QTLs. Second, effect size estimation error is greater for rare alleles, and when effect size is over-estimated, an association is more likely due to winner’s curse, leading to a negative correlation between effect size and minor allele frequency (28).

To avoid variation in power across allele frequency, we repeated the eQTL and aseQTL analysis, down-sampling our population to 50 individuals in each test, such that 40 individuals were drawn from the more common genotype and 10 individuals were drawn from the less common type for each SNP tested. As a consequence, for every SNP we test, sample sizes of major and minor genotype classes are equalized regardless of allele frequency in the population. We also measured effect sizes in this subsample to avoid any relationship between allele frequency and effect size estimation error. Despite reducing our sample size by half, we still detected 594 eQTLs and 670 aseQTLs, when using the most significantly associated SNP per gene (above a p-value threshold corresponding to FDR = 0.1; p<2.6×10^−5^ for eQTLs, p<8.2×10^−5^ for aseQTLs). In addition, we decoupled the identification of associations from the estimation of effect size by comparing allele frequencies of SNPs identified as eQTLs with these SNP’s effects on ASE, avoiding the double-testing issue responsible for winner’s curse.

Consistent with mutation-selection balance, an eQTL’s effect on ASE was negatively correlated with eQTL allele frequency (p < 0.05, correlation coefficient = −0.154, n=251) and total expression effect size was negatively correlated with aseQTL allele frequency (p<0.01, correlation coefficient = −0.104, n=670). eQTL and aseQTL allele frequency were also negatively correlated with the corresponding effect size when we designated a random significantly-associated SNP per gene as the focal eQTL or aseQTL (Figure S4 C,D). One possible explanation for the stronger association between eQTL allele frequency and ASE effect than between aseQTL allele frequency and total expression effect may be that ASE variation results from *cis* regulatory variation while total expression variation is determined by both *cis* and *trans* regulatory variation. Since we only map local QTLs that mainly act in *cis,* extra noise from *trans* regulatory variation likely contributes to total expression variation, weakening the association between allele frequency and total expression effect.

We also investigated the allele frequency spectra of the eQTLs and aseQTLs detected using the downsampling approach. In this case it was appropriate to use the frequencies of all SNPs tested as a neutral hypothesis because false positive and false negative rates are independent of minor allele frequency. The minor allele frequencies of the eQTLs and aseQTLs detected with downsampling were rarer than the frequencies of all SNPs tested (Figure 5). The skew towards rare alleles was stronger here than in the QTLs detected with the whole data set, perhaps because the reduced sample size of the downsampling approach allows us only to detect large effect QTLs, which are likely to be under stronger negative selection.

**Figure 5:**
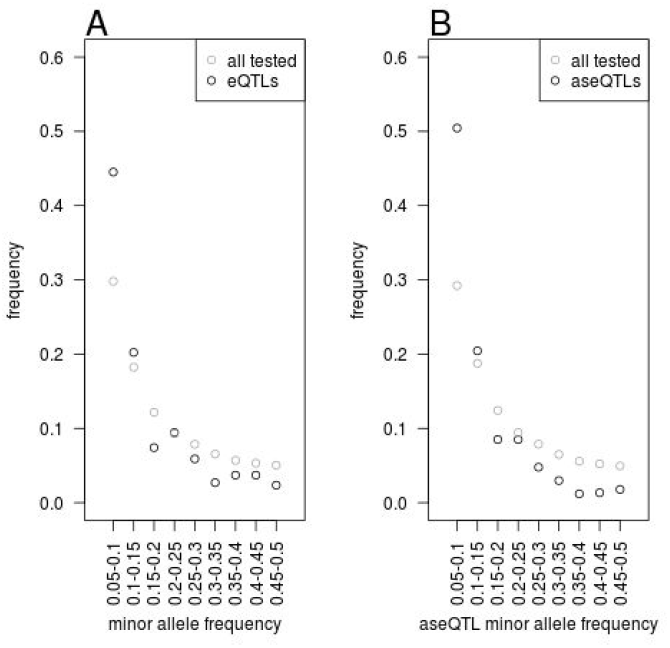
The site frequency spectra of QTLs detected in the frequency-controlled subsample.

It is important to note that some of our QTLs may not be causal alleles, but are instead in linkage disequilibrium with a causal allele. However, this is unlikely to strongly affect the allele frequencies of the QTLs we detect because the extent of linkage disequilibrium is constrained by similarities in allele frequency since the coefficient of linkage disequilibrium (D) is highest when the frequencies of both loci are similar. Consistent with this inference, power analyses have shown that a causal SNP and a tagging SNP in incomplete LD must have similar allele frequencies for a GWAS to successfully identify an association with the tagging SNP (29). Therefore, our conclusions about the allele frequencies of QTLs should be robust to the inclusion of non-causal linked alleles.

Our mapping of QTLs for expression and allele-specific expression genome-wide in a single population of *C. grandiflora* demonstrates that the frequencies and phenotypic effect sizes of these QTLs are consistent with mutation-selection balance. In addition, the enrichment of eQTLs in CNSs directly upstream of genes further supports CNS’s potential role as regulatory elements; however, the large number of QTLs discovered outside of conserved regions suggests significant turnover in regulatory elements between species. Alternatively, QTLs may create new deleterious regulatory interactions, instead of disrupting conserved functional sites. Taken together, our results, indicate that much of local expression variation observed at the population level is deleterious and support the role of mutation-selection balance in maintaining genetic variation within populations.

## Materials and Methods

### Study system and plant material

*Capsella grandiflora* is an obligately outcrossing member of the Brassicaceae family with a large effective population size (Ne~600,000), relatively low population structure and a range that spans northern Greece and southern Albania(26, 30). In June 2010, we collected seeds from approximately 400 plants growing in a roadside population of *C. grandiflora* near Monodendri, Greece (Population Cg-9(30)). We germinated and grew one individual from each parent in the University of Toronto greenhouses and performed crosses between independent random pairs of plants to generate the seeds used in this study. By growing the parents in a common environment and then assaying their progeny in a common environment, we reduced the influence of maternal effects and unknown micro-environmental effects on gene expression.

Approximately 10 seeds from each cross were sterilized in 10% bleach followed by 70% ethanol, placed on sterile plates filled with 0.8% agar with Mursashige-Skoog salts (2.15 g/L), stratified in the dark at 4^o^C for one week, and then allowed to germinate in a growth chamber at 22^o^C and 16 hour photoperiod. After one week, we transplanted two of the seedlings from each cross into 4 inch pots filled with ProMix BX soil and returned the pots to the growth chamber. After another week, pots were thinned down to one seed per cross. Throughout the experiment, pots were randomized once every week to minimize location effects.

Leaf tissue from young leaves was collected for RNA extraction four weeks after transplanting and immediately flash frozen in liquid nitrogen. RNA was extracted using plant RNA extraction kits (Sigma) from 2 or 3 samples from each plant. The extracted RNA was quantified with a Qubit spectrophotometer and the samples from each plant were pooled such that each pool contained the same amount of RNA from each sample. RNA was sequenced at the Genome Quebec Innovation Centre on two flow cells with 8 samples per lane. Reads were 100bp long and paired end. We extracted DNA from leaf tissue using a CTAB based protocol. Whole genome sequence from each individual was obtained through 100 cycles of paired-end sequencing in a Hiseq 2000 with Truseq libraries (Illumina), with three individuals sequenced per lane.

### Genomic data

We mapped DNA sequence data to the *C. rubella* reference genome(31) with Stampy v1.0.19. After bioinformatic processing with Picard tools, we realigned reads around putative indels with GATK RealignerTargetCreator and IndelRealigner and compressed the resulting bams with GATK ReduceReads. Raw SNP calls were generated by joint calling of all samples in GATK v2.81 UnifiedGenotyper. We subsequently followed GATK Best Practices for Variant Quality Recalibration using a high confidence subset of the raw calls generated by filtering snps for concordance with common variants (minor allele frequency > 0.11) in a species-wide sample of *C. grandiflora* (3) as well as suspect realignments (transposable elements, centromeres, 600bp intervals containing extreme Hardy-Weinberg deviations, 1kb intervals with evidence of 3 or more snps in reference-to-reference mapping). A relatedness analysis revealed that six individuals were more related to eachother than expected in an outcrossing population, perhaps because of introgression from *C. rubella,* so we removed these individuals from the analysis. We measured population structure using fastStructure on a set of 56,011 biallelic snps distributed genome wide that had been pruned for LD following the recommended analysis stream(27).

To map RNA reads, we constructed our own codon-only reference sequence by stitching together the exons and UTRs of each gene into a scaffold using reference gene annotations (31). We mapped to this codon-only reference using Stampy 1.0.21 with default settings. We chose to use Stampy over other RNA-specific aligners, like Tophat, because visual examination of alignments showed that Stampy was better at mapping reads containing multiple polymorphisms, reducing the potential for false associations between expression level and the genotypic variants that affect mapping (Fig. S6). RNAseq readmapping for two individuals was very poor quality (<10% reads mapped and paired correctly), so these individuals were removed. Our final sample size was 99 individuals.

Expression level was measured with the HTSeq.scripts.count feature of HTSeq, which counts the number of read pairs that map to each gene. We normalized the read counts of each sample for library size by dividing read counts by the median read count of the entire sample. Previous studies on human gene expression have found interactions between GC content, lane, and expression level (12), but we did not detect this (Fig. S7). Genes with a median expression level below five reads per individual before normalization were removed from the analysis, leaving a total of 18,692 genes.

### Mapping local eQTL

We selected SNPs for our eQTL analysis by finding all SNPs within the window spanning 5 kb upstream of the gene’s transcription start site and 5kb downstream from the gene’s transcription end site. We chose the 5kb range because a previous study in *Arabidopsis thaliana* mapping associations between expression and SNPs within 30kb of the gene found that 87% of local eQTLs were located within 5kb of the gene (23). SNPs were categorized as occurring in 0-fold degenerate sites, 4-fold degenerate sites, 2 or 3-fold degenerate sites, 5’UTRs, 3’UTRs, introns, stop codons, or intergenic regions based reference annotations (31). In addition, we identified SNPs located in non-coding sequence conserved across the Brassicaceae family (3). We only included SNPs with at least 10 heterozygous individuals and 10 individuals that were homozygous for the common allele in our sample.

We wrote set of Python scripts to test for associations between expression level and genotype at a nearby SNP. These scripts are available at https://github.com/emjosephs/eQTL. We mapped eQTLs by conducting a Mann-Whitney U test on the null hypothesis that gene expression does not differ between individuals that were homozygous for the common allele and individuals that were heterozygous. We used non-parametric statistics because expression data is not normally distributed. We used the Mann-Whitney U test function in SciPy (“scipy.stats.mannwhitneyu”), which uses a continuity correction and corrects for ties. 8,302 of our genes had ties in expression level between individuals and these ties on average involved 4.5 individuals(32). In addition,we compared common homozygotes to heterozygotes because we expect most local eQTLs to act in *cis* and thus be additive (12), and because not being limited by the sample size of rare homozygotes allowed us to map eQTL at rarer alleles.

To avoid a relationship between allele frequency and sample size, we conducted a second eQTL analysis where we subsampled 50 individuals for each SNP tested so that 40 individuals had the most common genotypic category (usually the homozygote) and 10 had the less common genotypic category (usually heterozygote) (14). We chose these thresholds because they retained most individuals while allowing us to still test 3,972,771 of the 4,098,832 SNPs originally tested for eQTLs (96.9%).

For both eQTL analyses, we controlled for multiple testing by using a false discovery rate approach (33) and only considered eQTLs to be associated with expression if that association had a p value corresponding to a false discovery rate of <0.1. To avoid being biased by detecting multiple SNPs linked to only one causal site, we only selected one eQTL per gene, picking the SNP with the lowest p value for association. However, to investigate whether choosing the most associated SNP biased our results, we also performed all analyses with eQTLs that were randomly chosen from the pool of SNPs significantly associated with expression (FDR = 0.1). We calculated the expression effect size of eQTLs by taking the absolute value of the difference between mean expression in the common homozygote and mean expression in the heterozygote.

### Mapping aseQTL

If local eQTLs act in *cis,* they should have allele-specific effects and individuals heterozygous for an eQTL will show a larger difference in expression between alleles than individuals homozygous for an eQTL. To take advantage of this second signature of expression variation, we developed a method to test for allele-specific expression QTL, or ‘aseQTL’ (similar approaches have been used in humans (14)). We quantified allele-specific expression at all heterozygous sites inferred from the genomic data. We used the count of reads mapped to each allele, taken from the ‘AD’ values in a VCF file constructed from the RNAseq data using GATK Unified Genotyper to calculate an allele-specific-expression measure (‘ASE) for each gene in each individual. Specifically, we calculated the mean of the the differences in allelic expression values at all heterozygous sites across a gene and divided this mean by median expression level of all genes in the individual to control for sequencing depth. While we expected that our measure of gene-wide ASE would be more accurate when we required multiple heterozygous sites per gene, doing so did not strongly alter the number of aseQTLs we found or their allele frequency distribution, so we only required one heterozygous site per gene to measure ASE (Fig. S8). We designated aseQTLs as the most associated SNP per gene that had higher ASE in heterozygotes for that SNP than in homozygotes for that SNP. However, we also performed all analyses designating aseQTLS as a SNP that was randomly sampled from the set of SNPs that were significantly associated with expression (FDR = 0.1) and had greater ASE in homozygotes for that SNP than heterozygotes.

ASE measures were not normally distributed, so we used a Mann-Whitney U test to test the null hypothesis that ASE did not differ between individuals that were heterozygous at a given SNP and individuals that were homozygous for either allele at that SNP. As before, we used the mannwhitneyu function in the SciPy package. 8,334 genes had at least one tie between individuals for ASE value and an average of 4 individuals were involved in ties within these genes. We only tested SNPs where we had 10 individuals that were both heterozygous at the SNP and had a heterozygous marker site in the gene and 10 individuals that were homozygous at the SNP and had a heterozygous marker site in the gene, allowing us to test for associations at 17,880 genes.

As in the eQTL analysis, we conducted a second aseQTL analysis where we subsampled 50 individuals for each SNP tested such that 40 had the most common genotypic category (usually the homozygote) and 10 had the less common genotypic category (usually heterozygote). This sample size allowed us to test 3,841,452 of the 3,966,364 SNPs originally tested for aseQTL (96.8%). For both sets of analyses, we conducted a false discovery rate analysis as described in the eQTL section and, selected all SNPS with a p value below the FDR threshold of 0.1, we chose the most significantly associated SNP per gene for further analysis, with the additional requirement that heterozygous individuals have higher ASE than homozygous individuals. We calculated ASE effect size for aseQTLs and eQTLs by taking the difference between mean ASE in homozygotes and mean ASE in heterozygotes. We only report ASE effects for eQTLs located outside the coding sequence of these genes they regulate.

Preferential mapping of reference alleles compared to alternative alleles could lead to spurious ASE. To evaluate the importance of this effect, we simulated all of the possible reads spanning each heterozygous site, containing either the reference allele or an alternate allele using scripts from Degner et al(34). There were up to 200 reads possible for each site, although reads near the start and end of genes had fewer reads covering them since we discarded all reads that were less than 100bp long. We mapped these reads with the same program and settings we used for the real data, with the exception that these reads were single-ended. Out of 2,365,590 SNPs in coding regions, 19,017 SNPs (<1%) had unequal numbers of reads mapping from each allele. 11,339 (60%) of these sites had more reads that mapped with the reference allele than with the alternative allele, suggesting that there is some reference bias. The 19,017 SNPs with evidence of mapping bias occurred in 3,059 genes. Removing these genes from the analysis did not qualitatively affect the minor allele frequency of aseQTLs (Fig. S9)

### Permutation analysis

Conducting millions of tests for genotype-expression associations with a relatively small (n=99) sample size exposes us to two potential sources of bias that correlate with the allele frequency of the SNPs we are testing. First, smaller sample sizes at low frequencies reduce power to detect associations. Second, smaller sample sizes at low frequencies increase our risk of false positives because expression data is non-normally distributed and outliers in a small sample will have a disproportionate effect on the mean(13). We found this second possibility especially concerning because it is not conservative with respect to our hypothesis that purifying selection will maintain eQTLs and aseQTLs at lower allele frequencies.

To ensure that our conclusions about allele frequencies were not due to false positives being more common at low allele frequencies, we compared the eQTLs and aseQTLs we found with those discovered using permuted data. We constructed permutations by randomly shuffling the assignments between genotype and expression values or allele-specific expression values for each gene. This strategy allows us to retain the allele frequencies and spatial distributions of the SNPs we are testing along with the distribution of expression and allele-specific expression values of each gene. Each permuted set was analyzed using the same methods as the real data, with one exception: instead of calculating a FDR for each permuted data set, we used the p-value cut offs from the real data to identify ‘false-positive’ eQTLs and aseQTLs in the permuted data. The frequency distributions of these ‘false-positive’ QTLs were used as a null distribution The permutation analyses do not directly control for site type, recombination rate, or other factors that could both bias a SNP towards being an eQTL/aseQTL and reduce allele frequency. To ensure that these effects did not drive our observations, we divided our eQTLs and aseQTLs from the real data and from permuted data into subsets. First, for site type, we selected the most strongly associated eQTL and aseQTL per gene that came from a the site-type of interest. Our site types were 5’UTRs, 3’UTRs, introns, intronic CNSs, exons (divided into 5 regions based on distance from start and end of the gene), and upstream and downstream CNS and nonconserved regions. For upstream and downstream regions, we divided sites into those within 1 kb of the TSS/TES and those that were 1 to 5 kb from the TSS/TES.

To control for recombination rate, we divided SNPs into those coming from high recombination regions (> 3.45 cM/mB) and low recombination regions (< 3.45 cM/mB) using recombination rate data calculated by using a genetic map made from a cross between *C. rubella* and *C. grandiflora* (35). To test for confounding effects due to gene conversion, we divided SNPs into those whose mutations could be due to gene conversion (A or T and C or G) and others (A and T or G and C).

## Acknowledgements

We thank Niroshini Epitawalage, Amanda Gorton, and Khaled Hazzouri for lab assistance, J. Paul Foxe for collection assistance, Wei Wang for computer assistance, and Aneil Agrawal, Graham Coop, Asher Cutter, Alan Moses, Adrian Platts, Tanja Slotte, and Robert Williamson for helpful suggestions. Thomas Bureau, Mathieu Blanchette, Daniel Schoen, Paul Harrison, Alan Moses, Adrian Platts, and Eef Harmsen contributed to the Value-directed Evolutionary Genomics Initiative (VEGI) grant (Genome Quebec/Genome Canada) which supported this work, along with an NSF Graduate Research Fellowship to EBJ (DGE-1048376), and NSERC Canada and CFI grants to JRS and SIW.

